# Drug Functional Site-Unknown Molecular Targets Exemplified by SARS-CoV-2 pseudoknot and ribosomal frameshifting-modulators

**DOI:** 10.64898/2026.07.02.735960

**Authors:** Ahmed Mohamed Ragab, Christopher Llynard D. Ortiz, Yu-Tong Huang, Jian-Zhou Wang, Yu-Han Wu, Jin-Der Wen, Lee-Wei Yang

## Abstract

The –1 Programmed Ribosomal Frameshifting (–1 PRF) signal of SARS-CoV-2, driven by a conserved three-stemmed RNA pseudoknot (PK), is indispensable for viral replication and represents a structurally stable yet underexplored therapeutic target. Unlike rapidly mutating viral proteins, this RNA element offers an opportunity for durable intervention but has historically been considered “undruggable”. We developed an integrative drug discovery and characterization pipeline that combines molecular docking, molecular dynamics simulations, and dual-luciferase assays to systematically identify and validate frameshifting–efficiency (Feff) modulators from FDA–approved compounds. To move beyond traditional similarity-based screening, we introduced a contact–distribution–matching method, which ranks candidate compounds by comparing their predicted RNA interaction fingerprints with those of reference modulators. This computational approach, paired with experimental validation, enabled us to expand the repertoire of Feff modulators and establish correlations between binding patterns and functional outcomes. To uncover the underlying mechanisms, we applied steered molecular dynamics simulations and single-molecule optical tweezers measurements, revealing that Feff-enhancing modulators preferentially stabilize the remote stem (stem 3) of the PK, promoting variety of intermediate force species with the beginning base pairs of stem 1 being refolded, even after those base pairs have been unwound by ribosome. The refold of the tips of the stem 1 in turn “push back” the ribosome on the slippery sequence to result in enhanced frameshifting. On the other hand, Feff-suppressing modulators rigidify early stem regions, increasing resistance to ribosomal progression and increase the drop-off rate, eventually leading to a reduced –1 frame to 0 frame ratio in translation. Together, these findings provide the first integrated demonstration of how small molecules can modulate –1 PRF by altering RNA PK folding dynamics. More broadly, our framework establishes a generalizable strategy for rationally targeting structured RNAs with repurposed drugs and offers new opportunities to expand the druggable genome to include noncoding RNA elements and other biomolecular targets lacking known functional sites.

## Introduction

Structure-based rational drug design/discovery has become a central paradigm in modern targeted therapy. However, the design becomes difficult when the exact locations of these important targets’ functional sites are unknown. In such cases, previous computational efforts have often begun by predicting candidate active, ligand-binding, or regulatory sites before designing corresponding therapeutics. Dynamics-based approaches prioritize residues coupled to collective motions, including catalytic residues enriched near global hinges or mechanically constrained regions (Yang and Bahar, 2005). Pocket-detection and solvent-mapping methods instead scan the entire structure for ligandable cavities and evolutionarily conserved hot spots, which can then guide docking, virtual screening, fragment growing, or free-energy refinement (Halgren, 2009; Le Guilloux et al., 2009; Ludlow et al., 2015). Complementary allosteric and cryptic-site methods expand the search to distal communication centers or transient pockets that may not be apparent in a static apo structure (Huang et al., 2013; Huang et al., 2019; Meller et al., 2023). Nevertheless, before a functional assay can be put in place, these computationally predicted sites or (ligand-bound) pockets observed by structural biology do not necessarily suggest their functional relevance and consequently the ligands bind these sites may or may not impact the function. In this scenario, functional site prediction, subsequent validation and pocket-specific drug screening/design are two separate steps.

SARS-CoV-2’s RNA pseudoknot, modulating the translation of essential NS enzymes to virus survival, has previously unreported functional site but whose frameshifting efficiency was shown to be modulated by small molecule compounds (Sun et al., 2021). Here, RNA pseudoknots (Chang et al., 2019) function as regulatory elements that control translation through programmed recoding mechanisms (Dinman, 2012; Brierley et al., 2006; Caliskan et al., 2014). One such mechanism, −1 programmed ribosomal frameshifting (−1 PRF), enables ribosomes to shift reading frame during translation, thereby expanding coding capacity and regulating protein stoichiometry (Jacks et al., 1988; Atkins and Gesteland, 2010; Plant and Dinman, 2005). −1 PRF is directed by a conserved three-component signal consisting of a slippery sequence, a short spacer, and a downstream stimulatory RNA structure, typically a pseudoknot (Brierley et al., 1992; Somogyi et al., 1993; Plant and Dinman, 2008). The pseudoknot impedes ribosomal translocation, generating tension that promotes slippage of tRNAs into the −1 frame (Qu et al., 2011; Wen et al., 2008). This process is tightly regulated, and its efficiency (Feff) determines the relative production of viral proteins, making it essential for viral replication (Dinman and Wickner 1992; Plant et al., 2008).

Coronaviruses, including SARS-CoV-2, rely on −1 PRF to control the expression of their replication machinery. The viral genome contains overlapping ORF1a and ORF1b, where successful frameshifting enables translation of ORF1b, encoding key enzymes such as the RNA-dependent RNA polymerase (Kim et al., 2020; Bhatt et al., 2021). Feff is finely tuned, and even modest perturbations disrupt viral protein stoichiometry and replication efficiency (Plant et al., 2010; Kelly et al., 2020). Importantly, the −1 PRF signal is highly conserved across coronaviruses, highlighting its potential as a therapeutic target (Kelly et al., 2020; Neupane et al., 2020). The SARS-CoV-2 −1 PRF signal is stimulated by a three-stemmed RNA pseudoknot whose structure has been resolved by cryo-electron microscopy (Bhatt et al., 2021). Despite detailed structural characterization, the mechanism by which the pseudoknot modulates frameshifting remains debated. Early models suggested that frameshifting efficiency correlates with the mechanical stability of the RNA structure (Chen et al., 2009), whereas subsequent single-molecule studies demonstrated that conformational plasticity, reflected by the ability to adopt alternative folding intermediates, is a more accurate predictor of Feff (Ritchie et al., 2012; Ritchie et al., 2014; Neupane et al., 2020). These findings suggest that −1 PRF is governed by a complex interplay between RNA structure, dynamics, and mechanical response.

Targeting the pseudoknot with small molecules has emerged as a promising antiviral strategy. Compounds such as MTDB and Merafloxacin have been shown to modulate −1 PRF and suppress viral replication in vitro (Park et al., 2011; Sun et al., 2021), and recent screening efforts have identified additional modulators among FDA-approved drugs (Munshi et al., 2022). However, RNA-targeted drug discovery remains challenging due to the absence of well-defined binding pockets. To address these challenges, we developed an integrated computational–experimental framework for the discovery and mechanistic characterization of −1 PRF modulators. First, we established SimFDA, a chemical similarity–based screening approach that efficiently identifies candidate modulators from FDA-approved drug libraries. While this approach enables rapid exploration of chemical space, it does not account for the spatial specificity of RNA–ligand interactions. To overcome this limitation, we introduced contact-distribution matching (CDM) (Figure 1), which characterizes ligands based on the distribution of their contacts across the pseudoknot. By comparing interaction patterns at nucleotide resolution, CDM provides a functional descriptor that links binding location to modulation of Feff.

**Figure 1.**
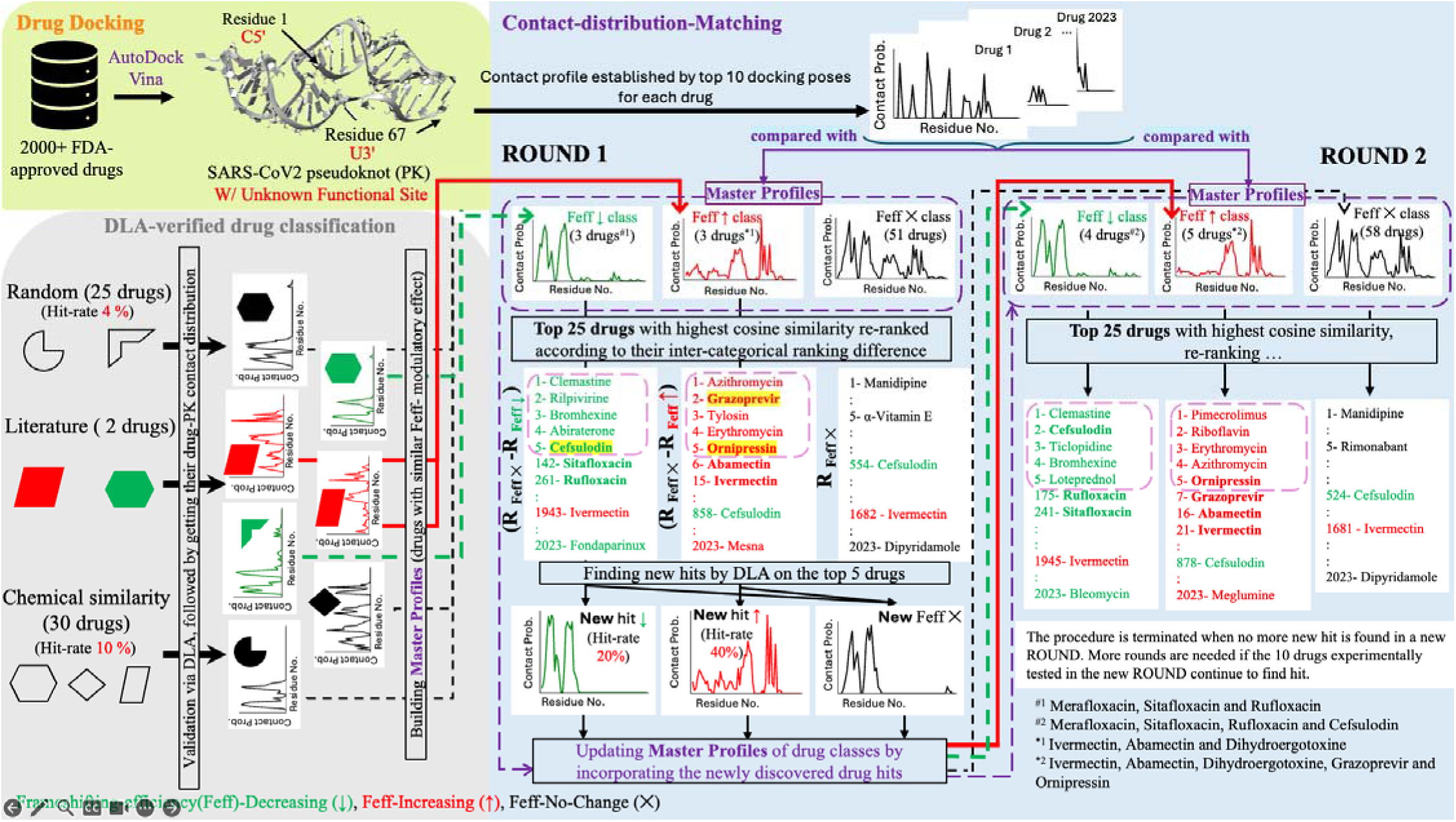
Overview of the Contact-Distribution-Matching (CDM) workflow for identifying small-molecule modulators of SARS-CoV-2 −1 PRF. FDA-approved compounds are docked to the SARS-CoV-2 frameshift-stimulatory pseudoknot (PK), and residue-level RNA–ligand contact probabilities are extracted from the top 10 docking poses to generate a 67-residue contact-distribution vector for each compound. Experimentally tested compounds are classified by dual-luciferase assay (DLA) according to their effects on frameshifting efficiency (Feff): Feff-decreasing compounds are shown in green, Feff-increasing compounds in red, and Feff-no-change compounds in black. Compounds with validated modulatory activity are used to construct class-specific reference, or “master contact profiles”. Candidate drugs are then compared with these master profiles using cosine similarity and ranked according to their similarity to the Feff-decreasing or Feff-increasing profiles, followed by an additional re-ranking procedure to consider the rank difference of the same drug in the Feff-no-change and in the Feff-decreasing/increasing categories, where the drug having a larger (Rank_Feff-no-change_ – Rank_Feff-decreasing/increasing_) values ranks higher. The top-ranked 5 candidates in the Feff-decreasing/increasing categories are experimentally tested by DLA in each ROUND, and newly validated hits, highlighted in yellow, are incorporated into the corresponding master profiles in the next ROUND to re-rank the contact distributions for the drugs, per aforementioned procedure. Drug names shown in **boldface** indicate compounds that were either included in the master profiles or subsequently identified as a new hit in the current ROUND through the CDM-guided selection process. This iterative screening strategy is repeated until no new hit is found in the top 5 in the Feff-decreasing/increasing categories, enabling progressive enrichment of compounds that d crease or increase SARS-CoV-2 −1 PRF efficiency.

We combined these computational approaches with dual-luciferase assays and single-molecule optical tweezers to quantify frameshifting efficiency and probe the mechanical properties of the pseudoknot. Optical tweezers measurements revealed that −1 PRF modulators alter the population of unfolding intermediates, supporting a central role for conformational dynamics in frameshifting regulation (Ritchie et al., 2012; Ritchie et al., 2014). To further elucidate the underlying mechanism, we performed molecular dynamics (MD) and steered molecular dynamics (SMD) simulations to characterize ligand-dependent conformational changes and unfolding pathways. These simulations demonstrate that ligand binding induces distinct alterations in the mechanical response of the pseudoknot, providing a direct link between RNA–ligand interactions and ribosome-induced force propagation.

Collectively, our results support a domain-specific mechanochemical model for −1 PRF modulation. Compounds that decrease Feff preferentially stabilize the proximal stems (Stems 1 and 2), thereby restricting conformational fluctuations required for efficient frameshifting. In contrast, compounds that increase Feff enhance flexibility and stabilize interactions in the distal region (Stem 3), promoting alternative conformations that facilitate ribosomal slippage. This spatially resolved mechanism reconciles previously conflicting models by demonstrating that the functional outcome of ligand binding depends on its location within the pseudoknot rather than on global stability alone. By integrating chemical similarity screening, interaction-pattern analysis, and mechanistic interrogation, this work establishes a unified framework linking chemical space, RNA dynamics, and translational regulation. These findings not only advance our understanding of −1 PRF but also provide a generalizable strategy for targeting structured RNA elements in viral genomes and beyond.

## Results

### Source of the known frameshifting efficiency (Feff) modulators

#### From literature

Sun et al., (2021) screened over 4,800 chemical compounds (including FDA-approved drugs) using a high-throughput approach and identified Merafloxacin and Ivermectin as a −1 programmed ribosomal frameshifting (−1 PRF) suppressor and enhancer, respectively. However, Ivermectin was later excluded from further consideration due to its high cytotoxicity. We independently verified these findings *in vitro* using dual-luciferase assay (DLA) (Grentzmann et al., 1998), confirming that Merafloxacin reduced frameshifting efficiency (F_eff_) by approximately 50%, while Ivermectin enhanced Feff by ∼20% at a concentration o 20 µM (Figure 2).

**Figure 2.**
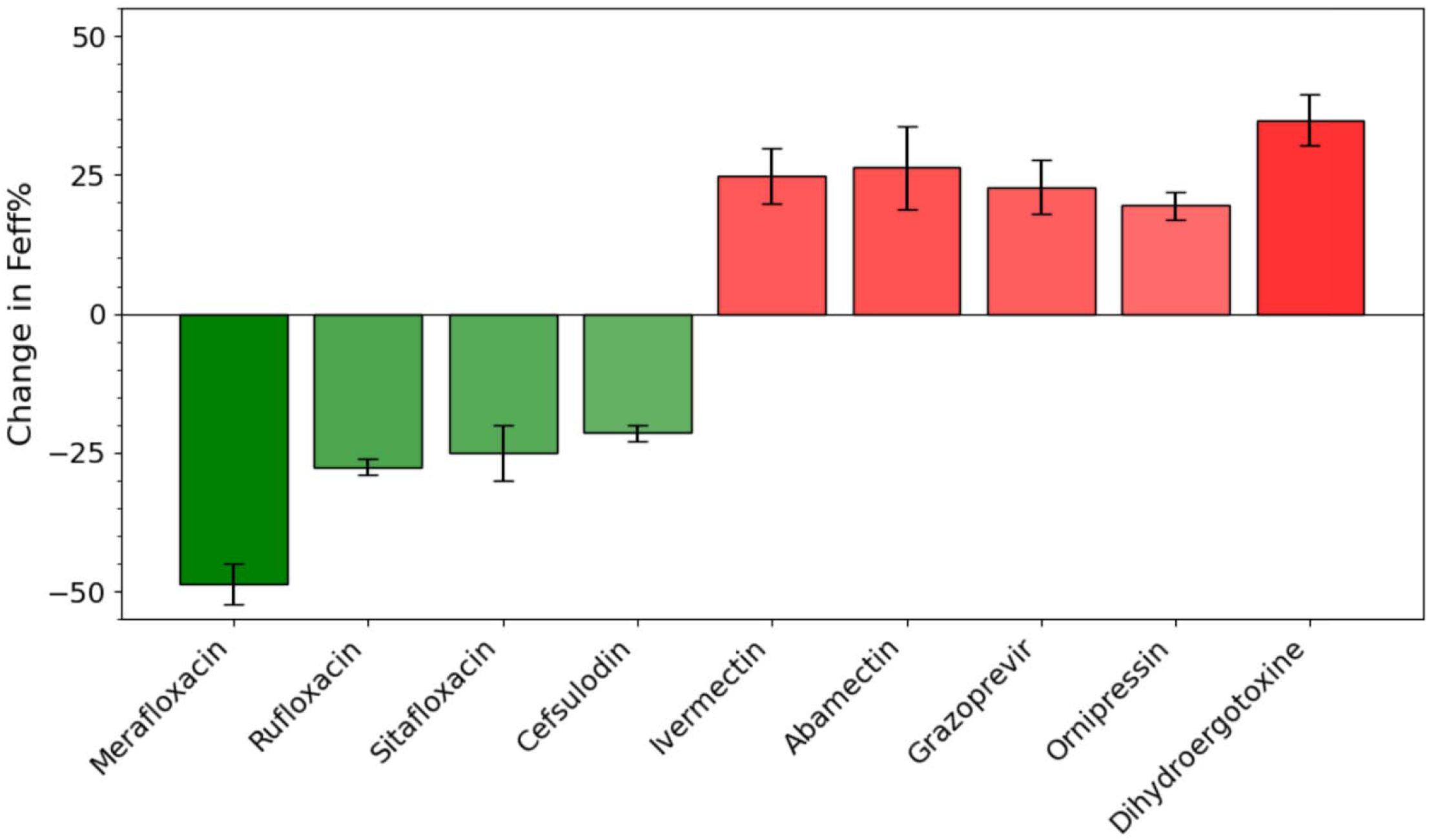
Repurposed FDA drugs modulate the frameshifting efficiency of SARS-CoV-2 PK. Change in frameshifting-efficiency (Feff) of SARS-CoV-2 –1 PRF in vitro in presence of hit drug candidates predicted by SimFDA. Rufloxacin, Sitafloxacin and Piromidic acid are analogous to Merafloxacin, while Abamectin and Moxidectin are analogous to Ivermectin. Dihydroergotoxine was identified from a random drug screening. Green bars represent suppression of –1 PRF by Feff-decreasing drugs, while red bars represent enhancement of –1 PRF by Feff-increasing drugs. Error bars represent standard error of the mean SEM from at least 5 replicates. Drugs are tested at the concentration of 20 uM.

#### Random search

To benchmark the predictive performance of this approach, we randomly screened 25 FDA-approved drugs from our in-house library using the DLA. From this unbiased screening, Dihydroergotoxine was identified as an Feff-increasing drug, enhancing Feff by ∼ 35% (Figure 2).

#### Chemical similarity search by an in-house engine, SimFDA

To expand the pool of potential modulators, we implemented **<u>SimFDA</u>** (https://dyn.life.nthu.edu.tw/SimFDA/), a chemical-similarity-based search engine comparing chemical fingerprints of the examined 2013 US FDA drugs, to identify FDA-approved drugs chemically similar to known Feff-modulators and rank-order them per their Tanimoto similarity to the query compounds in Sun’s paper (Sun et al., 2021) – Merafloxacin and Ivermectin. The search retrieved 25 Merafloxacin analogs and 2 Ivermectin analogs. When testing using the DLA, two Merafloxacin analogs – Rufloxacin and Sitafloxacin – were identified as Feff-decreasing drugs, lowering Feff by 28% and 25%, respectively (Figure 2). In contrast, Abamectin, analog of Ivermectin, enhanced Feff by ∼21%. Collectively, these results demonstrate that SimFDA achieved a high hit-rate in identifying novel Feff-modulatory compounds.).

### Contact-Distribution-Matching (CDM) identifies new −1 PRF modulators with high hitting rate

To identify novel modulators of the SARS-CoV-2 –1PRF signal, we implemented a **function-initiated approach** that prioritizes functional effectiveness over the traditional paradigm of structural site prediction. As illustrated in **Figure 1**, this process began with the experimental confirmation of initial –1PRF modulators – specifically the suppressor **Merafloxacin** and the enhancer **Ivermectin** – to establish baseline functional signals. By docking these compounds and the entire FDA-approved library to the cryo-EM-resolved pseudoknot (PK) structure, we extracted heavy-atom contact probabilities for each of the 67 nucleotides from the top 10 docking poses to construct unique **67-dimensional interaction vectors**. These vectors were aggregated into class-specific **“master contact profiles”** for –increasing, –decreasing, and –neutral categories, providing a spatial fingerprint of functional modulation.

The **Contact-Distribution-Matching (CDM)** workflow utilized these master profiles to rank candidate drugs from the library using **cosine similarity**. To maximize functional selectivity, we applied a **re-ranking procedure** that prioritized drugs exhibiting high similarity to active profiles while remaining distinct from the neutral profile, specifically by calculating the rank difference between categories. As new hits were validated through dual-luciferase assays (DLA), they were incorporated into the master profiles in subsequent rounds, iteratively refining the interaction signature required for –1PRF modulation.

Our results demonstrate that the CDM method is a significantly more robust tool for drug discovery compared to traditional paradigms. We evaluated the effectiveness of our approach by comparing **hit rates**, where a “hit” is defined as a drug changing the frameshifting efficiency (Feff) by 20% or more:

<u>Random Search</u>: Unbiased screening of FDA-approved drugs yielded a low hit rate of only 4% (1/25).

<u>Chemical Similarity (SimFDA)</u>: Searching for structural analogs of known modulators improved the hit rate to 10% (3/30), identifying compounds like Rufloxacin and Sitafloxacin.

<u>Contact-Distribution-Matching (CDM)</u>: By focusing on interaction patterns rather than chemical scaffolds, the CDM method achieved a superior hit rate between 20% and 40%.

This substantial increase in predictive power enabled the identification of chemically diverse modulators, such as the Feff-decreasing antibiotic **Cefsulodin** and the Feff-increasing antiviral **Grazoprevir**, which were not captured by structural similarity alone. These results confirm that CDM serves as a highly efficient platform for drugging complex RNA structures, effectively circumventing the challenges of targeting molecules with unknown functional sites

### Simultaneously finding functional sites after identifying functional modulators

After collecting all the DLA-validated hits, we are then able to plot the contact probability-coded PK structure as shown in Figure 3B, where the spatially concentrated colored probabilities mark the Feff-increasing (red) and Feff-decreasing (green) functional sites.

**Figure 3.**
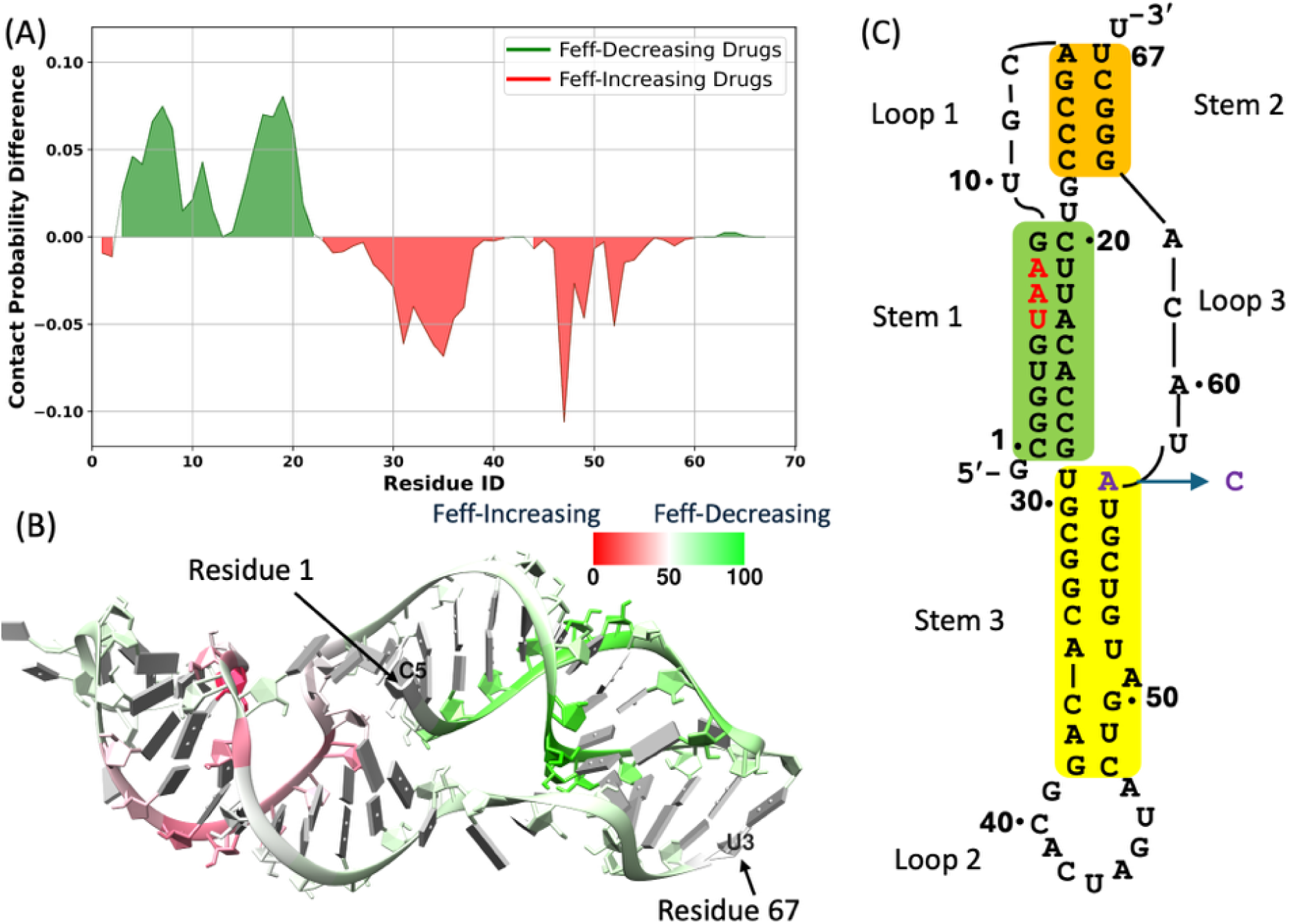
Differential interactions of Feff-increasing vs. Feff-decreasing compounds with the SARS-CoV-2 frameshift-stimulatory pseudoknot (PK). (A) Per-nucleotide difference in heavy-atom contact probability between the PK and the two compound classes. Green-colored nucleotides represent contacts enriched with Feff-decreasing drugs; red-colored nucleotides represent contacts enriched with Feff-increasing drugs. (B) Residue-specific differences from panel A mapped onto the cryo-EM three-Stem PK structure (Bhatt et al., 2021) by encoding values in the B-factor field after min–max normalization. Green residues (Stems 1 and 2) interact preferentially with Feff-decreasing drugs; red residues (Stem 3) interact preferentially with Feff-increasing drugs. The PK backbone is shown in grey, with arrows indicating the 5′ (Residue 1) and 3′ (Residue 67) ends. (C) Secondary-structure schematic of the SARS-CoV-2 −1 PRF signal. Stem 1 (green), Stem 2 (orange), and Stem 3 (yellow) form the three-Stemmed PK. The red UAA triplet in Stem 1 encodes an in-frame stop codon, blocking 0-frame translation unless −1 frameshifting occurs. The blue arrow marks the only sequence difference between SARS-CoV and SARS-CoV-2 (A→C).

**Figure 3.**
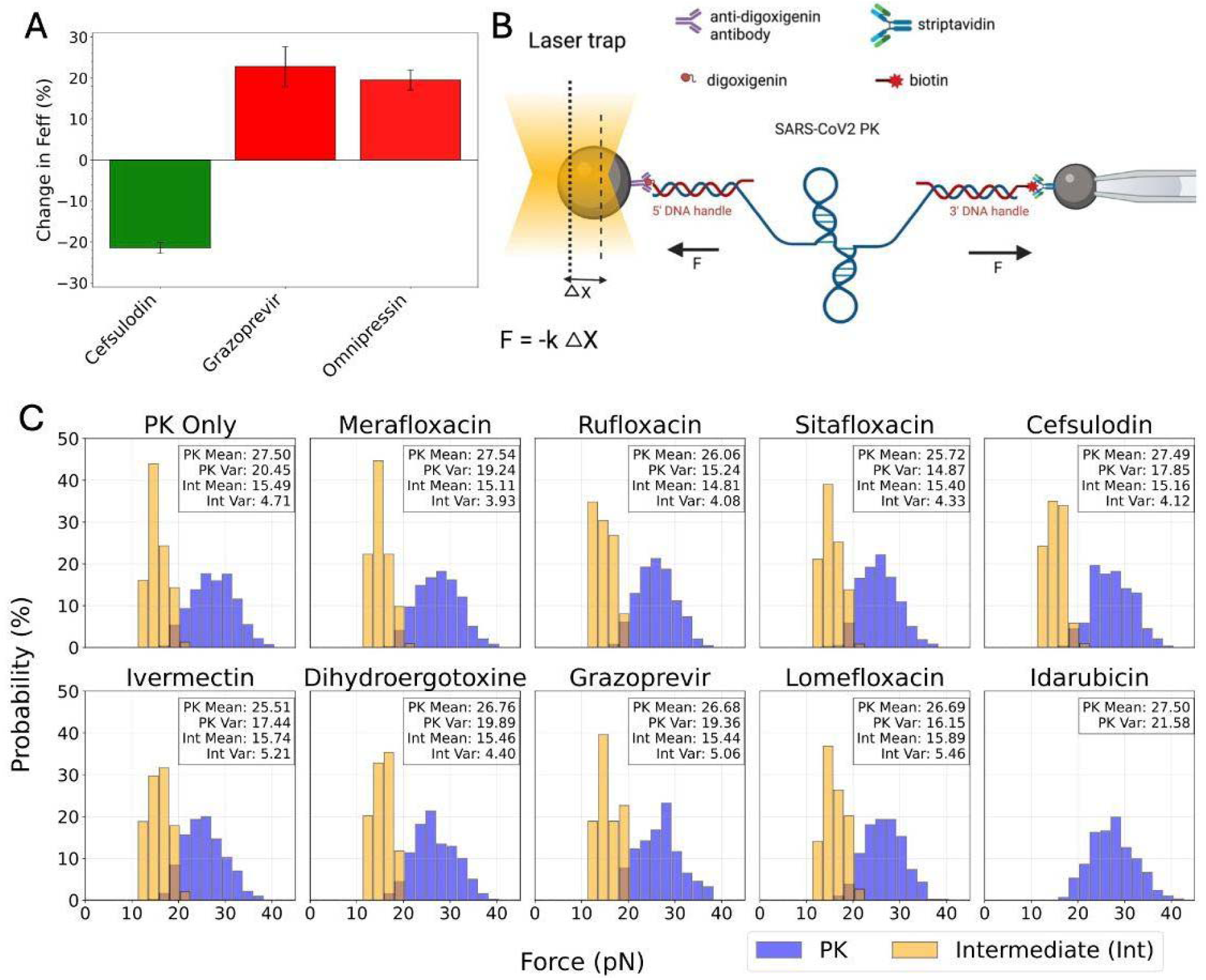
Drug-induced structural dynamics of the SARS-CoV-2 pseudoknot. (A) Change in frameshifting-efficiency (Feff) of SARS-CoV-2 –1 PRF for newly identified drug candidates predicted by Contact-Distribution-Matching method. Cefsulodin (green bar) is predicted to be a Feff-decreasing drug, while Grazoprevir and Ornipressin (red bars) are Feff-increasing drugs. Error bars represent SEM from at least 5 replicates. Drugs are tested at the concentration of 20 uM. (B) Schematic of single-molecule optical tweezers assay. The SARS-CoV-2 –1 PRF stimulatory pseudoknot (PK) is tethered between two beads through complementary DNA handles (biotin-streptavidin and digoxigenin-anti-digoxigenin). One bead is held in a laser trap while the other is fixed by a micropipette. As the trap is displaced, the restoring force (F) is measured from bead displacement (Δx) using Hooke’s law (F = – kΔx) to monitor unfolding transitions of the PK. (C) Unfolding-force distributions with and without drugs. Probability histograms for the mechanically strongest PK conformation (blue) and lower-force intermediates (orange) under different drug conditions: PK-only, Feff-decreasing (Merafloxacin, Rufloxacin, Sitafloxacin, and Cefsulodin), Feff-increasing (Ivermectin, Dihydroergotoxine, and Grazoprevir), Lomefloxacin is a non-changing-Feff analogous to Merafloxacin, while Idarubicin is a non-changing-Feff drug identified from random screening. Histograms show the probability distribution of force-induced unfolding events for the PK conformation (blue) and alternative intermediate states (orange).

### Drug-induced pseudoknot mechanical remodeling revealed by optical tweezers and Steered MD

To better understand the molecular basis of frameshifting modulation, we next investigated how Feff-modulatory drugs alter the mechanical stability and conformational landscape of the SARS-CoV-2 pseudoknot (PK). Using single-molecule optical tweezers (Figure 3B), we traced the changes in the PK unfolding forces and structural heterogeneity. Optical-tweezers measurements revealed distinct mechanical patterns. Feff-decreasing compounds such as Merafloxacin, Rufloxacin, Sitafloxacin, and Cefsulodin stabilized the folded PK, showing higher average unfolding forces and narrower distributions (Figure 3C). In contrast, Feff-increasing drugs, including Ivermectin, Dihydroergotoxine, and Grazoprevir, produced broader distributions and greater variance in intermediate force species, implying enhanced conformational plasticity (Figure 3C). Taken together, these data support a structural mechanism in which reduction in PK conformational plasticity corresponds to −1 PRF suppression, while increased flexibility promotes higher frameshifting efficiency.

To understand why these mechanical signatures differ, we wanted to analyze the per-nucleotide drug-PK contact patterns of the two drug classes. It worth mentioning that, in addition to the hit molecules identified from Contact-Distribution-Matching and SimFDA methods, we validated another Feff-decreasing FDA-approved drug, Nafamostat, reported by Munshi et al., (2022), and we found that it resulted in a 25% reduction in Feff (supplementary figure). By incorporating Nafamostat, we now have ten Feff-modulatory compounds in total: five Feff-decreasing (Merafloxacin, Sitafloxacin, Rufloxacin, Cefsulodin, and Nafamostat) and five Feff-increasing (Ivermectin, Abamectin, Dihydroergotoxin, Grazoprevir, and Ornipressin). Then, we generated aggregated contact profiles for each category by combining per-nucleotide contact probabilities across individual drug-PK interactions (supplementary figure).

We then computed the differential contact probability by subtracting the Feff-increasing drugs’ profile from that of the Feff-decreasing drugs at each of the 67 nucleotides (Figure 4A). This analysis revealed that Feff-decreasing drugs preferentially contact Stem 1, Loop 1, and portions of Stem 2. In contrast, Feff-increasing compounds predominantly engage with Stem 3, while also interacting with specific residues at Stem 1–Stem 3 junction (e.g., C1, G2, C27, G28) (Figure 4A-C). To visualize these nucleotide-level interaction preferences, we normalized the contact probability differences and mapped them onto the cryo-EM structure of the PK as B-factors in the PDB file. This structural representation highlights the distinct contact patterns of each drug class (Figure 4B).

**Figure 4.**
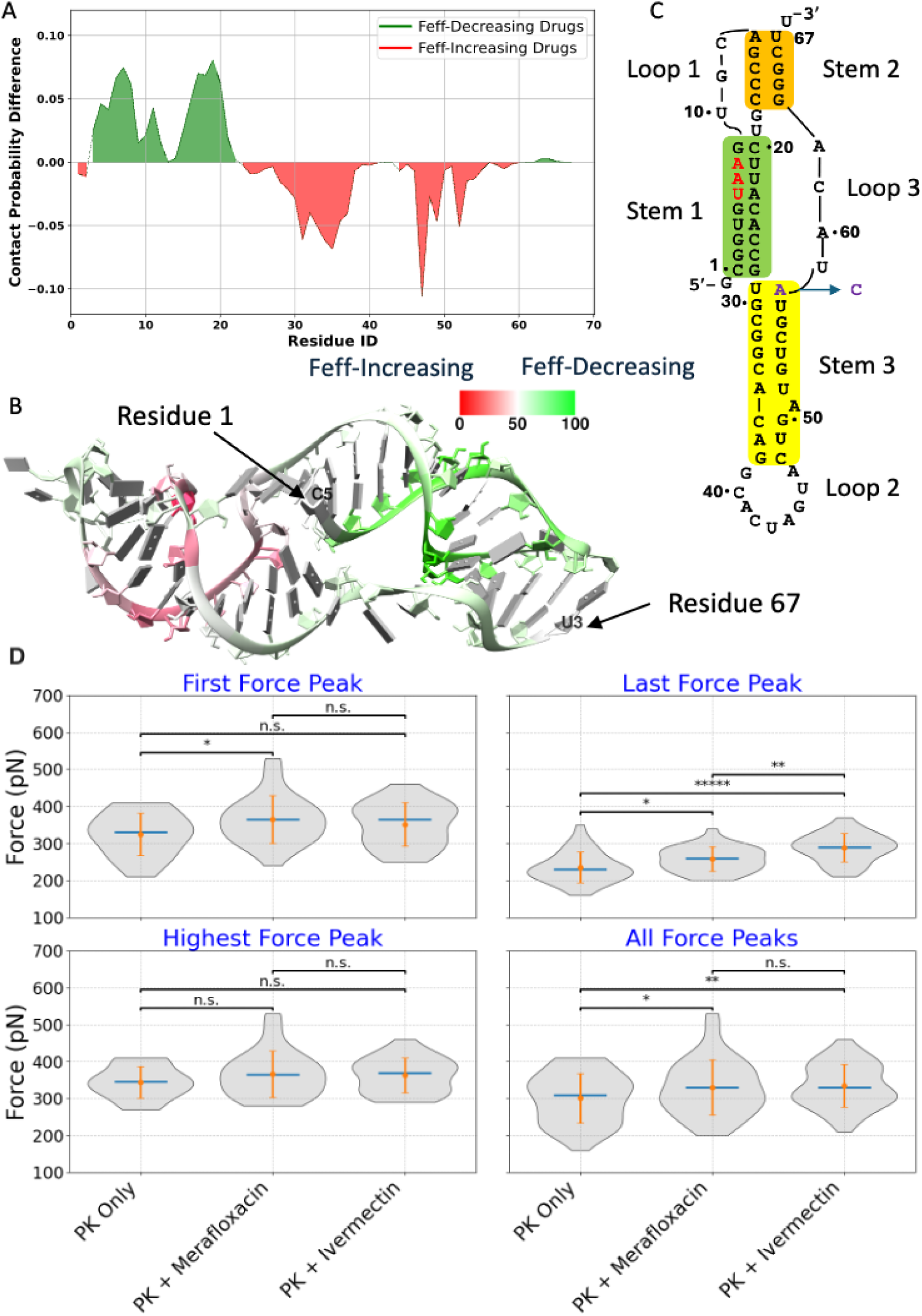
Differential interactions of Feff-increasing vs. Feff-decreasing compounds with the SARS-CoV-2 frameshift-stimulatory pseudoknot (PK). (A) Per-nucleotide difference in heavy-atom contact probability between the PK and the two compound classes. Green-colored nucleotides represent contacts enriched with Feff-decreasing drugs; red-colored nucleotides represent contacts enriched with Feff-increasing drugs. (B) Residue-specific differences from panel A mapped onto the cryo-EM three-Stem PK struct re (Bhatt et al., 2021) by encoding values in the B-factor field after min–max normalization. Green residues (Stems 1 and 2) interact preferentially with Feff-decreasing drugs; red residues (Stem 3) interact preferentially with Feff-increasing drugs. The PK backbone is shown in grey, with arrows indicating the 5′ (Residue 1) and 3′ (Residue 67) ends. (C) Secondary-structure schematic of t e SARS-CoV-2 −1 PRF signal. Stem 1 (green), Stem 2 (orange), and Stem 3 (yellow) form the three-Stemmed PK. The red UAA triplet in Stem 1 encodes an in-frame stop codon, blocking 0-frame translation unless −1 frameshifting occurs. The blue arrow marks the only sequence difference between SARS-CoV and SARS-CoV-2 (A→C). (D) Violin plots of rupture forces from 30 constant-velocity trajectories per system (PK-only, PK + Merafloxacin, PK + Ivermectin). Panels show the First, Last, and Highest force peaks per trajectory, and All Force Peaks (concatenated). Gray violins represent distributions; blue lines indicate medians; orange points with error bars indicate mean ± SD. Brackets indicate pairwise group comparisons (PK-only vs. PK + Merafloxacin; PK-only vs. PK + Ivermectin; PK + Merafloxacin vs. PK + Ivermectin) using two-sided Welch’s t-tests. Significance: * p < 0.05; ** p < 0.005; etc.,; n.s., not significant. Force values are reported in picoNewtons (pN).

We next asked how these binding preferences influence the pseudoknot’s unfolding pathway under force. To address this, we performed constant-velocity Steered Molecular Dynamics (SMD) simulations on three systems – PK-only, PK + Merafloxacin (Feff-decreasing), and PK + Ivermectin (Feff-increasing). Each system was simulated 30 times, where O5’ of the first nucleotide and O3’ of the last nucleotide of the RNA are pulled at a velocity of 10 m/s. From each trajectory, we extracted three representative rupture-force values: the first, highest, and last force peaks (Figure 4D) (Methods). Consistent with the optical-tweezers data, Merafloxacin markedly increased the first rupture force (mean ≈ 365 pN) compared with PK-only (≈ 324 pN), confirming stabilization of early unfolding intermediates. Ivermectin, in contrast, produced the highest last rupture forces (mean ≈ 289 pN versus 235 pN for PK-only), suggesting that it strengthens the terminal stem (Stem 3) and delays its unfolding.

To further evaluate how Feff-modulatory drugs influence the stability of specific PK domains, we used the X3DNA package (reference) to predict the number of base pairs throughout the simulation time and determine the unfolding kinetics of individual stems. In addition to Merafloxacin and Ivermectin as Feff-decreasing and Feff-increasing drug candidates, respectively, we further simulated other verified Feff-decreasing drugs such as (Sitafloxacin, Rufloxacin and Cefsulodin) and Feff-increasing drugs such as (Abamectin, Dihydroergotoxine (Dihydro.) and Ornipressin). We compared the unfolding trajectories of PK-only structures with those bound to Feff-increasing or Feff-decreasing drugs. Then, for each system, we calculated the time required for each Stem to be fully unfolded (i.e., reach zero base pairs). We observed that, on average, Feff-increasing drugs notably stabilized Stem 3, while slightly destabilizing Stem 1 (Figure 5B), compared to the “PK-only” and “Pk + Feff-Decreasing” systems. Conversely, the Feff-decreasing drugs did not show a statistically significant overall difference when compared to “PK-only”, except that Cefsulodin stabilized Stem 1 (longer unfolding time) and destabilized Stem 1 (shortened unfolding time) (Figure 5A).

**Figure 5.**
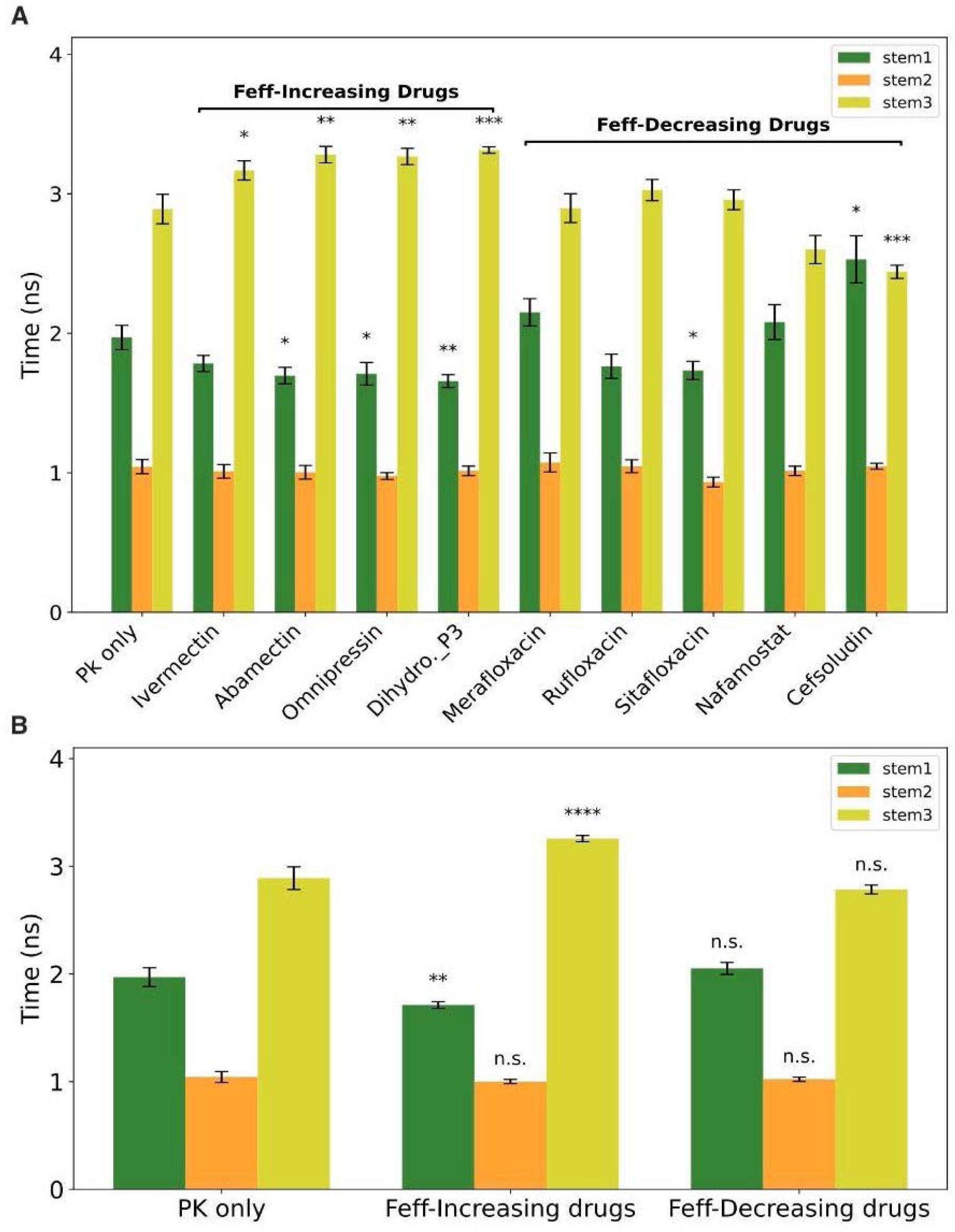
Constant-velocity SMD analysis of PK unfolding in the presence of Feff-modulating drugs. Bars show the average time (ns) for each PK’s stem (stem 1: green; stem 2: orange; stem 3: yellow) to reach zero base-pairing under constant-velocity pulling. (A) Individual drug-bound trajectories are plotted to the right of the PK-only control, while Feff-decreasing compounds (Merafloxacin, Rufloxacin, Sitafloxacin, Nafamostat and Cefsulodin) appear to the right most. Dihydro._p3 corresponds to docking pose 3, which engages the PK similar to other Feff-increasing drugs. (B) Mean unfolding times for PK-only, all Feff-increasing and all Feff-decreasing conditions. Error bars represent the standard error of the mean (SEM). Statistical significance versus PK-only is indicated by *p < 0.05, **p < 0.01, etc.; “n.s.” denotes non-significance.

To investigate how Feff-modulatory ligands influence PK stability upon applying pulling force, we performed constant-force SMD at ∼350 pN–matching the average force peak from constant-velocity runs (Fig. 1B). By pulling the O5′ and O3′ at termini nucleotides of PK in the three systems–PK alone, PK + Merafloxacin (an Feff-decreasing drug), and PK + Ivermectin (an Feff-increasing drug)–for 30 independent 1.6-ns trajectories. The time evolution of 5′–3′ distances differed across the three systems, indicating drug-dependent unfolding (Fig. 5A). By 0.8 ns, some replicas were fully extended while others remained partially unfolded, so we used this time point to compare intermediate ensembles. The 0.8-ns distance distributions revealed a stabilization hierarchy (Figure 6B): PK alone showed the greatest extension (mean ∼320 Å), Ivermectin shifted the ensemble toward more compact states (mean ∼250 Å), and Merafloxacin conferred the strongest resistance to unfolding (mean ∼180 Å). Consistently, intra-PK hydrogen-bond counts at 0.8 ns were highest with Merafloxacin, intermediate with Ivermectin, and lowest for PK alone. Collectively, the data support a drug-class-dependent stabilization hierarchy (Merafloxacin > Ivermectin > PK alone), consistent with Feff-decreasing ligands conferring greater mechanical resistance than Feff-increasing ligands.

**Figure 6.**
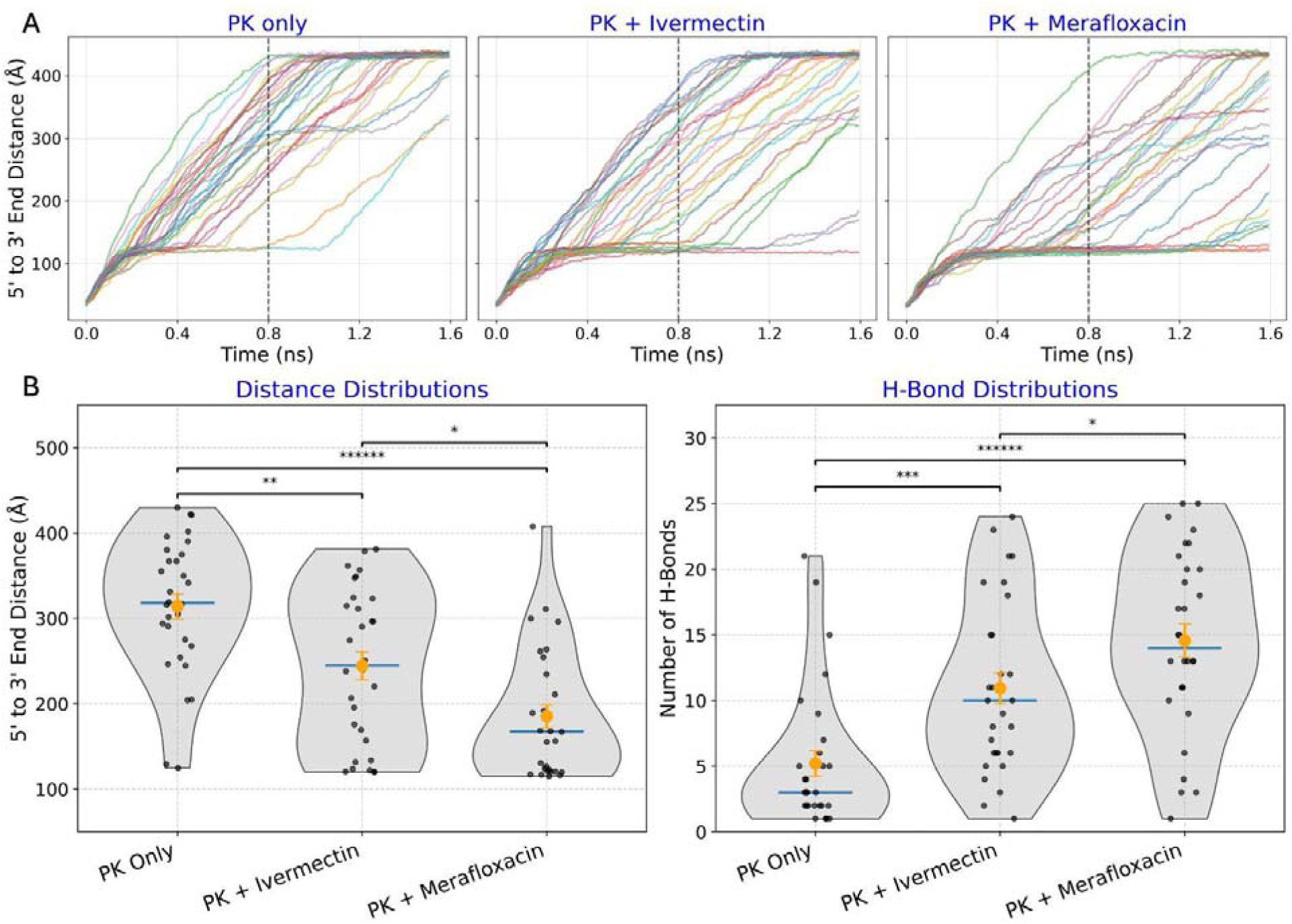
Constant-force SMD (350 pN) shows drug-class-dependent stabilization of the RNA pseudoknot (PK). (A) For each condition–PK-only, PK + Ivermectin (representative Feff-increasing drug), and PK + Merafloxacin (representative Feff-decreasing drug)–30 independent 1.6-ns trajectories are shown as 5′–3′ end-to-end distance traces (O5′ of the first nucleotide to O3′ of the last nucleotide). The vertical dashed line marks 0.8 ns, the time point used for ensemble analysis. (B) Distributions at ∼0.8 ns of (left) 5′–3′ distances and (right) intra-PK hydrogen bonds. Violin plots show the full distribution; black points (n=30) are individual trajectories; horizontal blue lines denote medians; orange markers indicate mean ± SEM. Pairwise comparisons are annotated with Welch’s t-test brackets (two-sided; Statistical significance versus PK-only is indicated by *p < 0.05, **p < 0.01, etc.; “n.s.” denotes non-significance if p ≥ 0.05).

We further analyzed the time it takes each stem of the three-stems-PK to reach zero base pairs for the three different systems. But as we noticed that some trajectories did not completely unfold till the end of simulation, we calculated the average time-to-zero base pairs only for those completely unfolded trajectories. At the constant-force of 350 pN, PK-only was the most readily unfolded system where all the three stems were almost fully unfolded, however, Ivermectin and Merafloxacin showed a significant stabilization of the different domains of PK, where the domains they contacted with took the longest time to be fully unfolded, and accounted for their resistance to unfolding. For instance, Ivermectin significantly increased the time to zero base pairs of stem 2 and 3 compared to PK-only, with only 83% of the trajectories becoming fully unfolded at stem 3. On the other hand, Merafloxacin showed a consistent increase of the global stability of the stems, achieving the largest average time needed to fully unfold the three stems in contrast to PK-only, and showing that stem 1 was remarkably stabilized than in the case of Ivermectin. We also noticed that the percentage of trajectories had fully unfolded three stems were ∼ 60% for stem 1, 83% for stem 2, and 70% for stem 3, respectively, clearly showing that Merafloxacin indeed leading to a global resistance of the PK’s domains for unfolding. We wanted to test if we may get a similar trend upon using a reasonably higher unfolding force that would indeed cause the PK structure for all the systems to be fully-unfolded. We performed constant-force SMD at 600 pN, a slightly higher than the highest force peak (∼ 540 pN) noticed in the constant-velocity analysis (Figure 4D)

By visually inspecting the constant-force SMD trajectories at 600 pN for the three systems, we noticed that indeed all the systems unfolded at the designated pulling force. Then, we performed the analysis of calculating the time-to-zero base pairs of each stem of the PK, and noticed that the unfolding time of stem 1 was a distinctive feature between the three systems. Ivermectin resulted in the shortest average time needed to first unfold stem 1 completely, while PK-only required slightly longer time to fully unfold the same domain, and Merafloxacin showed a significant increase in the time to unfold the same stem amongst the three systems (Figure 7B). In addition, stem 2 also took a significantly longer time to fully unfold in the case of Ivermectin and Merafloxacin when compared to PK-only, while there is no significant difference between the two systems with Feff-modulatory drugs. Similarly, stem 3 took a statistically significant longer time to completely unfold in case of Ivermectin and Merafloxacin in contrast to PK-only. These results further confirm that the contrasting preference of the drugs to specific domains of the PK indeed induces opposite effects in terms of PK unfolding dynamics.

**Figure 7.**
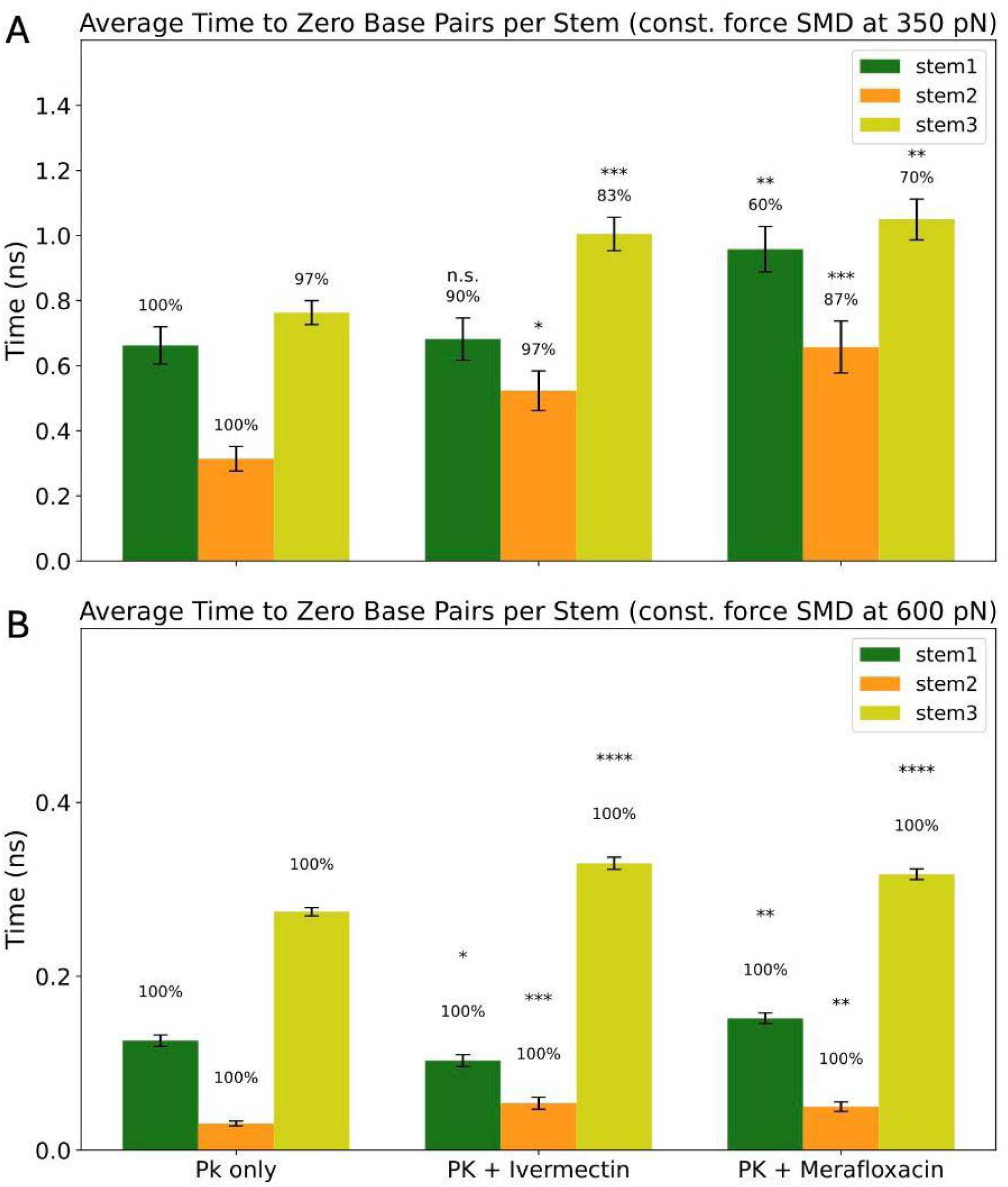
Constant-force SMD analysis of PK unfolding in the presence of Feff-modulating drugs at 350 and 600 pN. Bars show the average time (ns) for each PK stem (stem 1: green; stem 2: orange; stem 3: yellow) to reach zero base-pairing under constant-force pulling at 350 pN (A) and 600 pN (B). “PK-only” denotes the drug-free control; “PK + Ivermectin” represents a Feff-increasing condition; “PK + Merafloxacin” represents a Feff-decreasing condition. Error bars represent the standard error of the mean (SEM). Statistical significance versus PK-only is indicated by *p < 0.05, **p < 0.005, etc; “n.s.” denotes non-significance. Percent values above each bar represent the fraction of trajectories that fully unfolded within the simulation window.

We wanted to further study how the presence and absence of the Feff-modulatory drugs is affecting the refolding propensity of the PK. Using a similar protocol as that of the constant-force SMD, we allowed used the same three systems to unfold at a starting pulling force of 400 pN and simulated for 1 ns, then decreased the pulling force by 80 pN and simulated for 1 ns, progressively until it reach 0 pN, then allowed the system to simulated for 44 ns. This gradual reduction in pulling force allowed the fully unfolded RNA strand to restore base-pairing between its nucleotides and refold. We calculated the native base-pairing of the three stems, and noticed a very little propensity to restore the native base pairing pattern originally occupied by the PK (Figure 4C, add a supplementary figure showing the native contact of the three stems for the three systems). We were able to see that the refolding PK structures had a reasonable number of base pairing between its nucleotides (supplementary videos of representative refolding trajectories, and a figure of the total number of base pairs in the refolded RNA).

We looked in detail into those native base pairing the three systems were able to restore and found that C1G2--C27G28 nucleotides of stem 1 (check the nucleotides numbering in Figure 4C) were amongst the most frequently appearing base paired nucleotides. Interestingly, these nucleotides are showing a high contact probability by Ivermectin (Figure 4A), in contrast to Merafloxacin. We aimed to calculate the base pairing of these four nucleotides throughout the refolding simulation trajectories of the three systems after the first 6 ns. We calculated the AUC of these data and found that Ivermectin was showing the highest AUC value amongst the three systems, confirming that it indeed contributed to restoring the contact between these nucleotides. We wanted to further confirm it.

### Stem 3 stabilizer (Feff enhancer) promote the tip base pairs refolding in Stem 1 that further promotes –1 frameshifting

The pseudoknot was first unfolded using constant-force SMD by applying an initial force of 400 pN for 1 ns, followed by sequential reductions of 80 pN every 1 ns until the force reached 0 pN. The structures were subsequently simulated for an additional 44 ns at zero force, yielding a total simulation time of 50 ns. Although restoration of native base pairing across all three stems was generally limited, we observed pronounced differences in the recovery of specific structural elements located at the junction between stem 1 and stem 3.

Among these elements, the proximal stem-1 base pairs, C1–G28 and G2–C27, were of particular interest because they were frequently contacted by Feff-increasing compounds during docking and molecular dynamics simulations (Figure 3A-C). Analysis of the refolding trajectories revealed that Ivermectin substantially enhanced restoration of these base pairs relative to both the PK-only and Merafloxacin-bound systems (Figure 8A). Because these nucleotides are positioned at the stem 1–stem 3 junction and represent one of the earliest regions expected to be disrupted during ribosome-mediated unwinding, their recovery may serve as an indicator of productive pseudoknot remodeling rather than simple structural stabilization.

**Figure 8.**
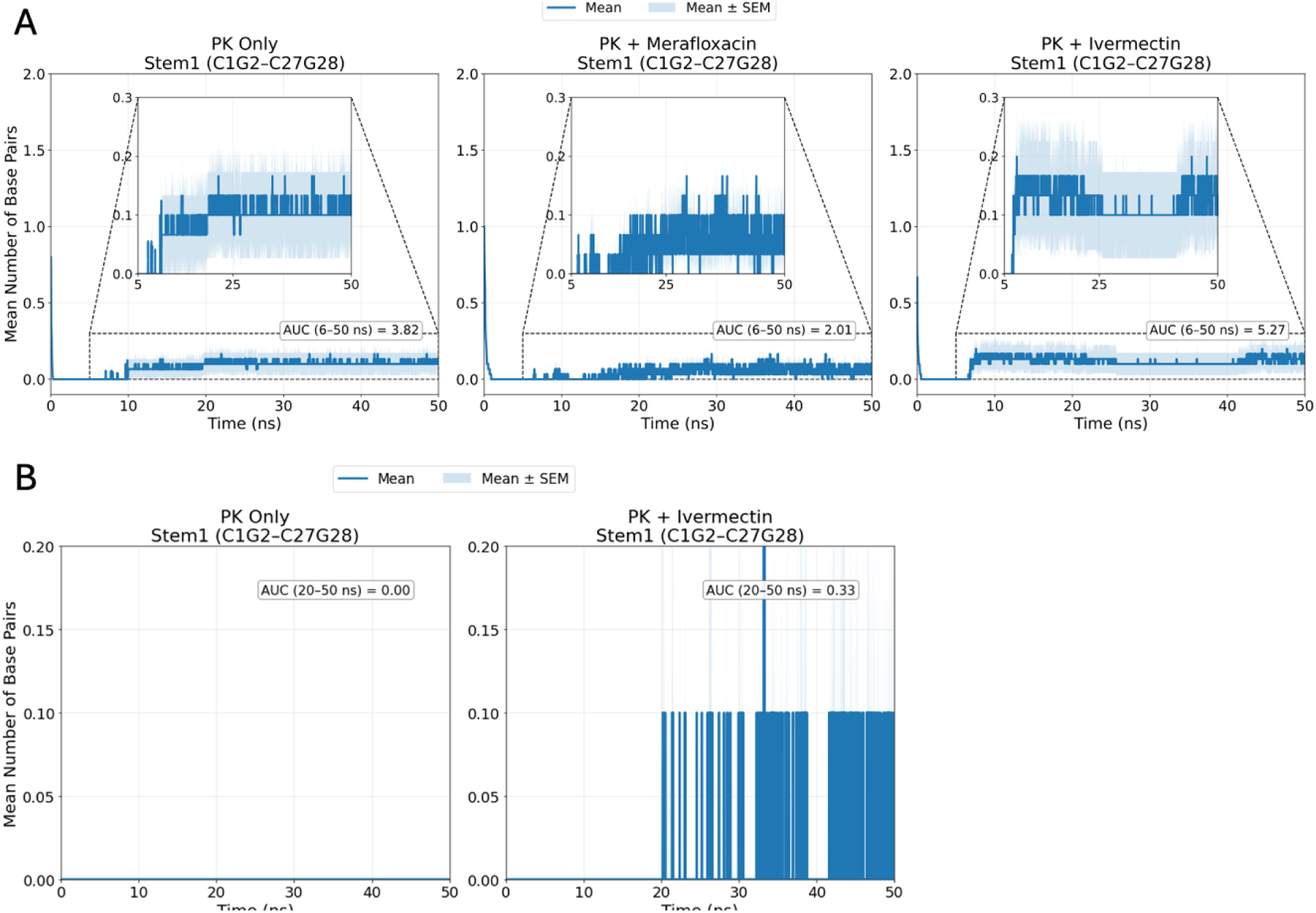
Differential recovery of proximal nucleotides of stem-1 of PK in presence and absence of Feff-modulatory drugs. **(A)** Recovery of proximal stem-1 base pairs during force-relaxation simulations. PK-only, PK + Merafloxacin, and PK + Ivermectin systems were simulated for 50 ns following constant-force SMD unfolding and stepwise reduction of the applied force to 0 pN. The number of restored proximal stem-1 base pairs, C1–G28 and G2–C27, was calculated over time and averaged across 30 independent simulations. Solid lines represent the mean, and shaded regions indicate the standard error of the mean (SEM). Insets show an enlarged view of the low-occupancy region (0–0.3 base pairs). The area under the mean recovery curve (AUC) was calculated from 6–50 ns to quantify cumulative restoration after completion of force relaxation. **(B)** Refolding of a partially unfolded pseudoknot intermediate at 0 pN. A conformation with unfolded stems 1 and 2 and intact stem 3 was simulated for 50 ns in PK-only and PK + Ivermectin systems. The number of restored proximal stem-1 base pairs, C1–G28 and G2–C27, was calculated over time and averaged across 10 independent simulations. Solid lines represent the mean, and shaded regions indicate SEM. AUC values were calculated from 20–50 ns.

To further evaluate this observation, we selected conformations in which stems 1 and 2 were completely unfolded while stem 3 remained intact. This structural state was chosen to mimic a ribosome-engaged pseudoknot in which upstream regions have been unwound while downstream structural elements remain folded. The selected conformation was simulated either in the presence of Ivermectin or after removal of the ligand to generate a corresponding PK-only system. Ten independent simulations were performed for each condition.

Remarkably, Ivermectin consistently promoted recovery of the proximal stem-1 base pairs C1G2–C27G28 after approximately 20 ns of simulation at zero force, whereas the PK-only system showed little or no tendency to restore these interactions throughout the simulation period (Figure 8B). These findings suggest that Ivermectin facilitates remodeling of partially unfolded pseudoknot intermediates into conformations capable of re-establishing critical stem-1 interactions. Rather than merely stabilizing the native structure, the compound appears to bias the folding landscape toward productive refolding pathways.

The observed behavior closely resembles the mechanism proposed by Hsu et al. (2021) for the highly efficient DU177 frameshift-stimulatory pseudoknot. In that study, optical tweezers, single-molecule FRET and molecular dynamics analyses demonstrated that frameshifting efficiency is strongly influenced by the ability pseudoknot to remodel folding intermediates rather than by structural stability alone. Specifically, of the partial unwinding of stem 1 by the ribosome relieves constraints on stem 2, allowing completion of pseudoknot folding and restoration of stem-1 interactions. This remodeled conformation is capable of retrieving nucleotides transiently occupied by the ribosome, thereby generating tension within the mRNA channel and promoting backward ribosomal slippage into the −1 reading frame.

Our observations suggest a similar mechanism for the SARS-CoV-2 pseudoknot. By stabilizing stem 3 and facilitating recovery of C1G2–C27G28, Ivermectin may promote formation of a refolding intermediate analogous to that described for DU177. Such intermediates may possess an enhanced ability to restore stem 1 interactions during translation, generating the mechanical tension necessary to stimulate −1 programmed ribosomal frameshifting.

Additional support for this model was obtained from the dual-luciferase assay results. Comparison of Feff-increasing and Feff-decreasing compounds revealed distinct effects on reporter expression (Figure 9). Feff-decreasing compounds produced significantly higher Renilla luciferase signals relative to Feff-increasing compounds, whereas Feff-increasing compounds yielded elevated Firefly luciferase expression. Given that Renilla luciferase is translated in the zero frame and Firefly luciferase is expressed only following successful −1 frameshifting, these results indicate that the two classes of compounds modulate translation through distinct mechanisms.

**Figure 9.**
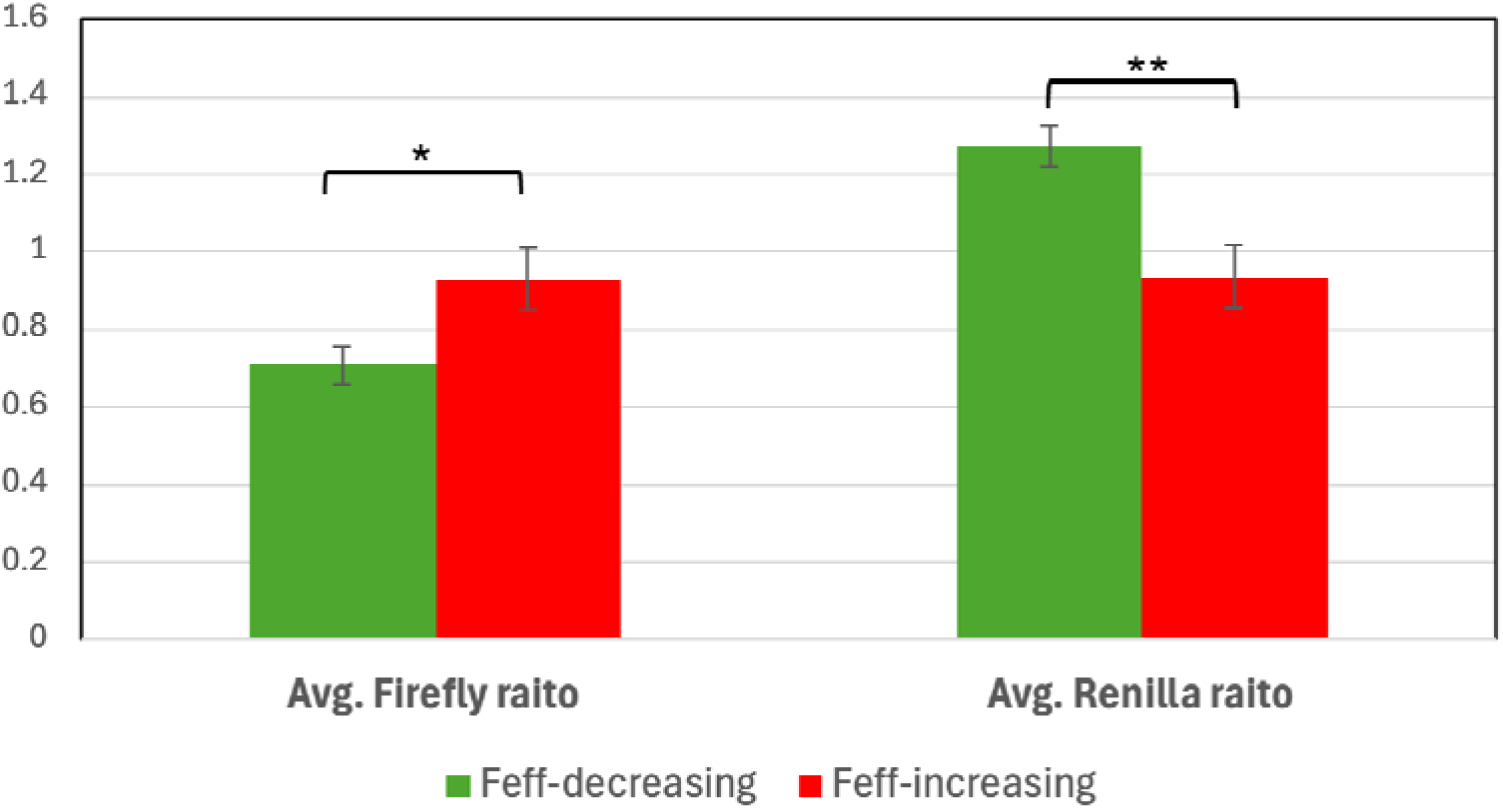
Feff-modulatory drugs’ effect on dual-luciferase reporter expression in vitro. Average relative Firefly and Renilla luciferase activities measured from SARS-CoV-2 dual-luciferase frameshifting constructs in the presence of Feff-increasing and Feff-decreasing compounds. Luciferase activities were normalized to those obtained from the corresponding control construct and averaged within each category based on the direction of frameshifting modulation. Bars represent the mean, and error bars indicate SEM. Statistical significance was determined using Student’s t-test, where (*) denotes P < 0.05 and (**) denotes P < 0.005.

Analysis of drug-binding preferences provides a structural explanation for these observations. Feff-decreasing compounds predominantly interacted with stem 1, whereas Feff-increasing compounds exhibited higher contact probabilities with stem 3 and the proximal region of stem 1. This distinction was consistently observed across docking and SMD simulations. Importantly, Feff-decreasing compounds also increased resistance of the pseudoknot to mechanical unfolding, resulting in a more rigid and constrained structure. In contrast, Feff-increasing compounds promoted the formation of intermediate-force species and enhanced recovery of critical stem 1 interactions during refolding.

These findings suggest fundamentally different effects on the pseudoknot folding landscape. Binding to stem 1 may rigidify early folding intermediates and restrict the structural rearrangements necessary for pr ductive pseudoknot remodeling. Such excessive stabilization may impede ribosomal progression, increasing the probability of ribosome stalling, premature termination, or ribosome dissociation from the mRNA. Consequently, repeated translation reinitiation events would generate increased amounts of Renilla luciferase relative to Firefly luciferase, consistent with the experimental observations (Figure 10).

**Figure 10.**
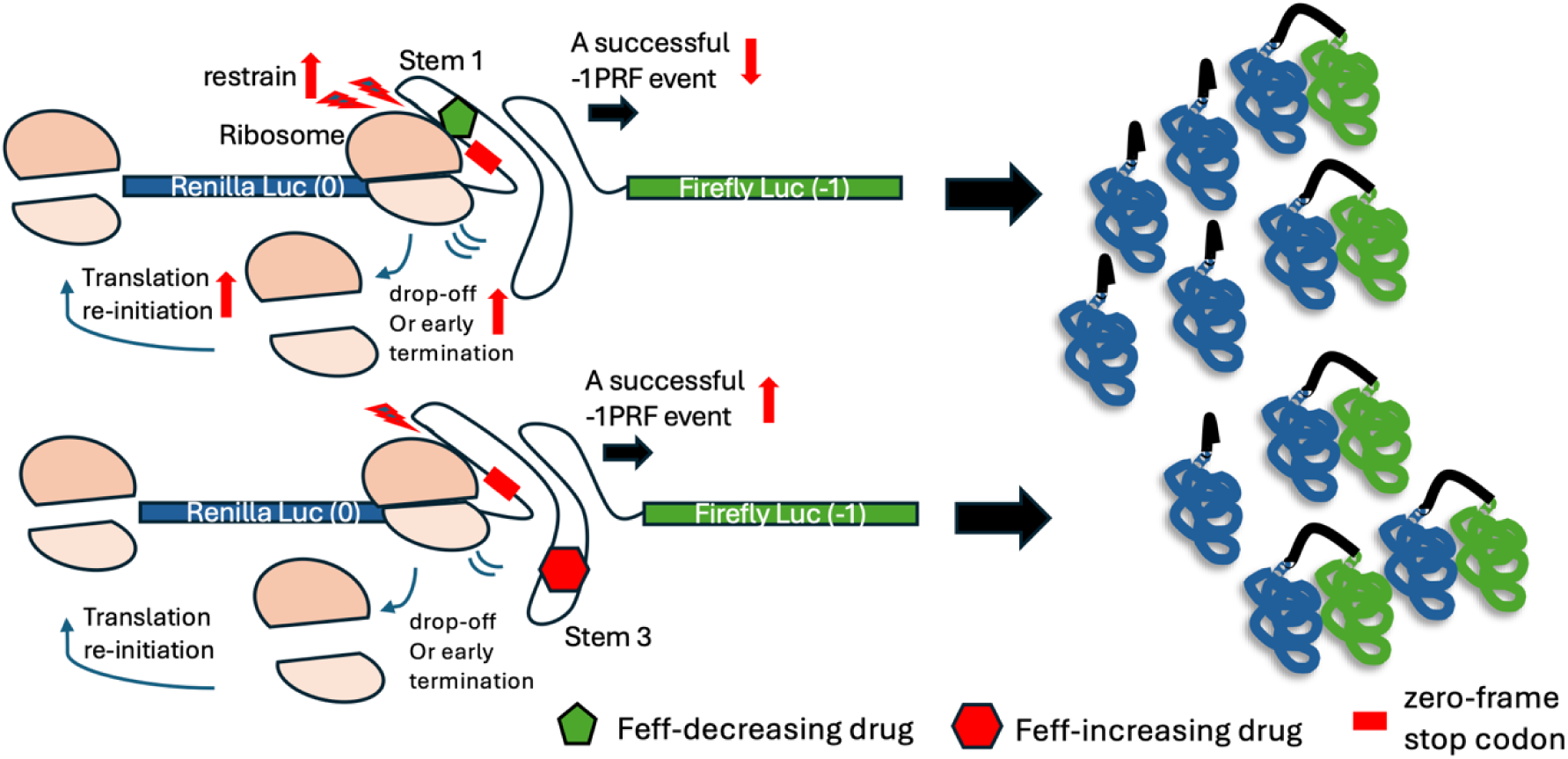
Schematic representation of the proposed effects of Feff-modulatory drugs on the SARS-CoV-2 −1 PRF dual-luciferase reporter system. The reporter mRNA contains Renilla luciferase (Luc) in the 0 frame, followed by the SARS-CoV-2 −1 PRF signal and Firefly luciferase in the −1 frame. The upper panel shows a Feff-decreasing drug bound near stem 1 of the pseudoknot. The lower panel shows a Feff-increasing drug bound near stem 3 and the proximal stem-1 region. Renilla and Firefly luciferase products are illustrated according to the indicated translation outcomes.

Conversely, compounds that preferentially interact with stem 3 appear to promote a distinct structural pathway. Although the ribosome remains capable of partially unfolding the upstream portion of stem 1, ligand-mediated stabilization of stem 3 facilitates remodeling of the partially unfolded intermediate into a more stable refolded conformation. This remodeled state may recover stem-1 interactions that were temporarily disrupted by the ribosome, thereby increasing tension along the mRNA and enhancing the probability of tRNA slippage into the −1 frame. As a consequence, successful frameshifting events occur more frequently, resulting in elevated Firefly luciferase expression (Figures 9 and 10).

Taken together, our results support a model in which frameshifting efficiency is governed not solely by pseudoknot stability but by the dynamic balance between unfolding and productive refolding. Feff-increasing compounds appear to enhance frameshifting by promoting remodeling of partially unfolded pseudoknot intermediates into tension-generating conformations capable of restoring critical stem-1 interactions. In contrast, Feff-decreasing compounds may over-stabilize stem-1-associated conformations and restrict the structural flexibility required for productive remodeling. This model is consistent with the mechanistic framework established for the DU177 pseudoknot (Hsu et al., 2021) and suggests that small molecules can modulate −1 programmed ribosomal frameshifting by reshaping the folding landscape of the stimulatory RNA structure rather than merely altering its thermodynamic stability (Figure 10).

## Discussion

Our study identifies a domain-specific mechanism by which small molecules modulate −1 programmed ribosomal frameshifting (−1 PRF) in SARS-CoV-2 through localized tuning of RNA pseudoknot (PK) structural plasticity. SimFDA enabled rapid identification of structurally related analogs of validated Feff modulators, leveraging chemical similarity for scaffold-based exploration (Figure 2). In parallel, CDM extended this search by matching RNA-ligand contact profiles across thousands of FDA-approved compounds to reference signatures of known Feff modulators, ranking candidates based on their domain-specific contact similarity (Figure 1). This dual strategy allowed us to identify both chemical analogs and mechanistically analogous but chemically diverse compounds – yielding multiple validated hits with high predictive success.

Furthermore, normalized nucleotide-level of the contact probability between the identified Feff-modulating small-molecules and the SARS-CoV2 PK showed that Feff-decreasing drugs such as Merafloxacin, Rufloxacin, Sitafloxacin, and Cefsulodin preferentially engage proximal domains such as Stems 1 and 2. Converse y, Feff-increasing compounds such as Ivermectin, Dihydroergotoxine, and Grazoprevir bind the distal Stem 3 and the Stem 1–Stem 3 junction (Figure 3A–C). These domain-specific contact preferences exerted noticeable changes to the mechanical unfolding of the PK in the presence of those drugs. Single-molecule optical tweezers experiments confirmed that Feff-decreasing ligands reduce structural heterogeneity, producing narrower unfolding-force distributions. In contrast, Feff-increasing compounds broaden the range of mechanical states (Figure 3B). This supports a mechanochemical model in which rigidification of basal stems inhibits frameshifting, whereas flexibility in distal stems facilitates it.

Steered molecular dynamics (SMD) simulations reinforced these contrasting domain-specific mechanical patterns. For instance, and in the presence of Merafloxacin, as a representative of Feff-decreasing drugs, the pseudoknot unfolds in fewer numbers, with a remarkably shortened 5’-to-3’ end distance of the trajectories and elevated average of H-bonding between the ligand and PK nucleotides (Figure 6A–B). With further inspection, this stabilization of the PK in presence of Merafloxacin seems to be attributed to the specific contact of Merafloxacin to Stem 1 mainly and Stem 2 with less extent (Figure 4B). Conversely, Feff-increasing drugs stabilize Stem 3, while destabilize Stem 1 (Figure 5A–B), increase the number of intermediate states, and delay final unfolding (Figure 7A–B). These trends were robust across both constant-force and constant-velocity SMD regimes, further establishing a causal link between site-specific flexibility tuning and frameshifting modulation.

The observed pharmacological effects parallel those of host cellular proteins known to regulate –1 PRF. ZAP-S, which suppresses frameshifting, binds to Stem 2, while SLBP, which enhances −1 PRF and viral replication, engages Stem 3 (Zimmer et al., 2021; Chen et al., 2025). The fact that drug-induced behaviors mirror these protein functions underscores a common mechanistic principle: local structural plasticity within the PK dictates frameshifting outcomes, whether modulated by small molecules or cellular cofactors.

These findings also reconcile long-standing models of −1 PRF regulation. Early studies proposed that frameshifting efficiency was directly proportional to the mechanical resistance of the pseudoknot – acting as a physical barrier to ribosome movement (Chen et al., 2009; Matsumoto et al., 2018; Namy et al., 2006; Kontos et al., 2001). However, subsequent single-molecule experiments argued instead for a structural plasticity model, wherein conformational heterogeneity – not rigidity – better predicts frameshifting efficiency (Ritchie et al., 2012; Halma et al., 2019). Our data resolve this dichotomy by showing that both models apply, depending on the structural domain engaged. Stabilization of basal stems suppresses frameshifting via reduced flexibility, whereas distal stem plasticity promotes slippage, consistent with the interactive pseudoknot refolding model proposed by Hsu et al. (2021).

In conclusion, we demonstrate that domain-specific modulation of RNA flexibility governs the frameshifting efficiency of the SARS-CoV-2 pseudoknot. Our data reveal a unified mechanochemical framework in which ligand binding dictates local unfolding dynamics, translating into functional control of −1 PRF. This framework not only resolves historical mechanistic debates but also provides a blueprint for rational RNA-targeted therapeutic development, applicable across viral systems that employ programmed ribosomal frameshifting.

